# The landscape of accessible chromatin in quiescent and post-myocardial infarction cardiac fibroblasts

**DOI:** 10.1101/2021.03.03.433814

**Authors:** Chaoyang Li, Jiangwen Sun, Qianglin Liu, Sanjeeva Dodlapati, Hao Ming, Leshan Wang, Yuxia Li, Rui Li, Zongliang Jiang, Joseph Francis, Xing Fu

## Abstract

After myocardial infarction, quiescent cardiac fibroblasts are activated and undergo multiple proliferation and differentiation events, which contribute to the extracellular matrix remodeling of the infarcted myocardium. We recently found that cardiac fibroblasts of different differentiation states had distinct expression profiles closely related to their functions. Gene expression is directly regulated by chromatin state. However, the role of chromatin reorganization in the drastic gene expression changes during post-MI differentiation of cardiac fibroblast has not been revealed. In this study, the gene expression profiling and genome-wide mapping of accessible chromatin in mouse cardiac fibroblasts isolated from uninjured hearts and the infarcts at different time points were performed by RNA sequencing (RNA-seq) and the assay for transposase-accessible chromatin with high-throughput sequencing (ATAC-seq), respectively. ATAC-seq peaks were highly enriched in the promoter area and distal areas where enhancers might be located. A positive correlation was identified between the transcription level and promoter accessibility for many dynamically expressed genes. In addition, it was found that DNA methylation may contribute to the post-MI chromatin remodeling and gene expression in cardiac fibroblasts. Integrated analysis of ATAC-seq and RNA-seq datasets also identified transcription factors that possibly contributed to the differential gene expression between cardiac fibroblasts of different states.

## Introduction

In response to myocardial infarction (MI), cardiac fibroblasts proliferate and differentiate into myofibroblasts expressing elevated levels of contractile proteins and extracellular matrix (ECM) proteins which are important in maintaining the structural integrity of the infarcted myocardium (1–3), but may have a detrimental effect on the long-term cardiac function due to permanent replacement fibrosis. Depletion of myofibroblasts after MI increased the early incidence of cardiac rupture (2) but also improved the long-term cardiac function (2, 3). Using lineage-tracing, we recently found that the proliferative state and myofibroblast state of cardiac fibroblasts were rather transient and largely limited to the first week after MI (1). Myofibroblasts then further differentiate into matrifibrocytes, a newly identified cardiac fibroblast differentiation state featured by the expression of some cartilage and bone genes and is likely an adaptation to the highly fibrotic environment. Cryoinjury-induced depletion of matrifibrocytes resulted in reduced scar stability and cardiac function, suggesting a critical role of matrifibrocytes. Using Affymetrix microarray, we recently reported the gene expression profiles of cardiac fibroblasts of different differentiation states and identified state-specific gene expression in these cells that corresponded to their unique functions (1). However, the mechanism responsible for the differential gene expression is still not known. Despite of a large amount of effort that has been made to explore the mechanism responsible for cardiac fibroblast activation and myofibroblast differentiation, our understanding of these processes is still relatively limited. An increasing number of studies have reported the presence of multiple signaling pathways involved in the post-injury activation and myofibroblast differentiation of CFs (4) besides the well-known TGFβ signaling (5–8), suggesting the complexity of these processes. Moreover, the molecular mechanism regulating the newly identified matrifibrocyte differentiation remains unknown. The accessibility of chromatin directly regulates gene transcription (9, 10). Thus, chromatin remodeling likely contributes to the dynamic gene expression during the sequential activation and differentiation events in cardiac fibroblasts after MI.

Using an optimized transposase-accessible chromatin with high-throughput sequencing (ATAC-seq) protocol modified for low cell numbers and a cardiac fibroblast lineage-tracing mouse line, we performed open chromatin profiling using cardiac fibroblasts from uninjured hearts and infarcts at different time points after MI. The same population of cardiac fibroblasts was also subjected to RNA sequencing (RNA-seq) analysis. The integrated analysis of ATAC-seq and RNA-seq data revealed the important role of chromatin remodeling in post-MI gene expression regulation in cardiac fibroblasts. Transcription factor (TF) binding motif analysis of the ATAC-seq data also revealed TFs that were possibly responsible for the dynamic chromatin accessibility and differential gene expression between cardiac fibroblasts of different differentiation states.

## Method

### Mice

Mouse lines used in this study are as follows: *Tcf21*-MerCreMer (11) and *R26*-EGFP (12). These 2 mouse lines were crossed to obtain *Tcf21*^*MCM/+*^;*R26*^*eGFP*^ mice.

### Animal procedures

To induce the activity of the MerCreMer, mice were treated with tamoxifen (Sigma T5648) dissolved in corn oil through intraperitoneal (IP) injections at a dosage of 75 mg/kg body weight/day for 5 days. MI was induced in mice via permanent surgical ligation of the left coronary artery (13). Briefly, mice were anesthetized using isoflurane and a left lateral thoracotomy was performed. The left coronary artery was identified and ligated just below the left atrium. For pain management related to surgical procedures, mice were given Carprofen at 5 mg/kg before the procedure and 12 hours after the procedure.

### Tissue process for imaging

Mouse heart samples were fixed in 4% paraformaldehyde for 3.5 hours, rinse in PBS for 30 min, and immersed in PBS containing 30% sucrose overnight at 4°C. The left ventricles of uninjured hearts and infarcts were carefully removed and trimmed for whole-mount imaging using a Leica SP8 confocal microscope.

### FACS

*FACS*. Heart tissue was digested as previously described with modification (14). Briefly, heart tissue samples were minced and digested in DMEM containing 0.75U/ml Collagenase D (Roche 11088866001), 1.0U/ml Dispase II (Roche 10165859001), and 1 mM CaCl_2_ at 37°C for 40 min. The slurry was then passed through a 100-μm cell strainer and then a 40-μm cell strainer. Cells were collected by centrifugation at 350 × g for 10 min. The cell pellet was then resuspended in PBS. *Tcf21* lineage-traced eGFP^+^ cardiac fibroblasts were sorted on FACSaria II (BD Biosciences). Gates were made based on WT eGFP^-^ control. Sorted eGFP^+^ cardiac fibroblasts were used for RNA-seq and ATAC-seq analyses.

### RNA-seq library construction and sequencing

Total RNA was extracted from 10,000 FACS-sorted *Tcf21* lineage-traced fibroblasts using the miRNeasy Micro Kit (217084) from Qiagen. cDNA libraries were constructed using the NEBNext single cell/low input RNA library prep kit for Illumina (E6420) from New England BioLabs. 150 bp paired-end sequencing was performed on the Illumina Hi-seq platform. Around 30 million read pairs were obtained for each sample. At least 2 samples of each group were analyzed.

### ATAC-seq library construction and sequencing

The ATAC-seq was performed following a protocol as previously described (15). Briefly, 10,000 FACS-sorted *Tcf21* lineage-traced fibroblasts were lysed to isolated the nuclei. Isolated nuclei were then incubated with the Tn5 transposase (TDE1, Illumina) and tagmentation buffer at 37°C for 30 minutes with shaking on a thermomixer at 500 g. Tagmentated DNA was purified using MinElute Reaction Cleanup Kit (Qiagen). PCR was performed to amplify the ATAC-seq libraries using Illumina TrueSeq primers and multiplex by indexes primers. After the PCR reaction, libraries were purified with the 1.1X AMPure beads (Beckman). The concentration of the sequencing libraries was determined by using Qubit dsDNA HS Assay Kit (Life Technologies). The size of the sequencing libraries was determined by means of High Sensitivity D5000 Assay with a Tapestation 4200 system (Agilent). 150 bp paired-end sequencing was performed on the Illumina Hi-seq platform. Around 50 million read pairs were obtained for each sample. 2 samples of each group were analyzed.

### RNA-seq data processing

RNA-seq reads that passed filters were trimmed to remove low-quality reads and adaptors by TrimGalore-0.6. The quality of reads after filtering was assessed by fastQC, followed by alignment to the mouse genome (MM10) by STAR (2.5.3a) with default parameters. Individual mapped reads were adjusted to provide TPM (transcripts per million) values with mouse genome as reference. Differential gene expression analysis was done using DESeq2 (16) and OneStopRNAseq (17).

### ATAC-seq data processing

Sequencing reads of all samples underwent adapter removal using TrimGalore-0.6, followed by quality assessment using FastQC. Reads were then aligned to the mouse reference genome MM10 using Bowtie 2.3 with the following options: – very-sensitive −X 2000 – no-mixed – no-discordant. Only unique alignments within each sample were retained in subsequent analysis. Moreover, alignments resulting from PCR duplicates or located in mitochondria were excluded. The mouse genome was tiled with consecutive non-overlapping 300bp bins. The accessibility of each of the 300bp bins was assessed by the number of fragments per million mapped (FPM) that was aligned to the bin. There was a total of 1,328,402 bins with at least one aligned fragment in at least one sample. Pairwise Poisson distance between samples was calculated using package PoiClaClu in R. The Principal Component Analysis (PCA) was also done in R.

### Peak calling and enrichment of genomic features in peaks

ATAC-seq peaks were called separately for each sample by MACS2 (18) with the following options: – keep-dup all – nolambda – nomodel. Peaks in individual samples from each group were subsequently merged using bedtools (https://bedtools.readthedocs.io/en/latest/). The annotations of genomic features, including transcription start sites (TSS), transcription end sites (TES), exons, introns, and CpG islands and repeat-elements: long interspersed nuclear elements (LINEs), short interspersed nuclear elements (SINEs), long terminal repeats (LTRs) and simple sequence repeats (Simple) were downloaded from UCSC genome browser. Promoters were defined as 500bp up- and downstream from the TSS of each annotated gene (TSS ± 500bp). Intergenic regions were defined as genomic regions before the TSS of the first gene and after the TES of the last gene in each chromosome, and in-between the TES and TSS of two consecutive genes. Peaks that did not overlap with annotated promoters were deemed as distal peaks. To evaluate the enrichment of the above genomic features with identified ATAC-seq peaks, a set of random peaks was first generated by matching the length of ATAC-seq peaks. The enrichment was then assessed by the ratio (or the log of the ratio) between the numbers of ATAC-seq and random peaks that overlapped with the corresponding genetic feature. The enrichment of transcriptional factor motifs in peaks was evaluated using HOMER (http://homer.ucsd.edu/homer/motif/).

### Assessment of promoter accessibility

Promoter accessibility in cardiac fibroblasts of each differentiation state was assessed by the number of ATAC-seq fragments (FPM) mapped to the defined promoter region (i.e.,TSS ± 500bp) in all samples from that stage. Z-score for each promoter at each stage was obtained by standardizing the FPM values stage-wisely, having mean 0 and variance 1 within each sample group. Promoters with z-scores below zero at all the stages were deemed to have constantly low accessibility; while those with z-scores above zero at all the stages were deemed to have constantly high accessibility. The remaining promoters were deemed to have dynamic accessibility. Pearson correlation was calculated between promoter accessibility and corresponding gene expression across stages. Promoters with accessibility that highly correlates with corresponding gene expression were then identified (with cor ≥ 0.5).

### Assessment of promoter CpG density

CpG density of annotated promoters (TSS ± 500bp) was assessed by function CpGDensityByRegion in R package BSgenome. In our analysis, when categorization of the promoter CpG density was needed, it was done as follows: low (<25), medium (≥25 and ≤75) and high (>75).

### Gene ontology and pathway analysis

Gene ontology (GO) enrichment analysis was performed using OneStopRNAseq (17).

### Gene regulatory network reconstruction

For each annotated gene, we first identified TFs with a binding motif that appears in ATAC-seq peaks overlapping with the regulatory region of the gene, which is defined as 100kb up- and down-stream of its TSS. A set of trans-regulators was then created for each gene to include all the identified TFs, and subsequently refined by retaining only TFs that were co-expressed with the gene (the absolute value of Pearson correlation ≥ 0.6). The network topological analysis and visualization were done in Cytoscape (19) with additional plugins: NetAnalyzer (20) and yFiles (21).

## Result

### Transcriptome profiling of cardiac fibroblasts of different differentiation state

To study the gene expression profiles of cardiac fibroblasts in different differentiation states, we employed the same CF lineage-tracing mouse line (*Tcf21*^*MCM/+*^*;R26*^*eGFP*^) that we used in our previous publication (**Figure 1A**) (1). *Tcf21*^*MCM/+*^*;R26*^*eGFP*^ mice were treated with tamoxifen and subjected to surgical induction of MI (**Figure 1B**). Whole-mount confocal imaging of the uninjured heart and the infarcted myocardium showed a significant expansion of the CF population after MI (**Figure 1C**). CFs were also enlarged after MI (**Figure 1C**), likely due to the highly abundant smooth muscle alpha-actin (αSMA) stress fibers. Moreover, CFs appeared to be more organized as the infarct became more stabilized, likely due to progressed infarct remodeling (**Figure 1C**). Freshly sorted *Tcf21* lineage-traced cardiac fibroblasts from the left ventricle of uninjured hearts (quiescent cardiac fibroblast) and infarcted myocardia at day 3 (proliferative early myofibroblast), day 7 (mature myofibroblast), 14 (transitioning matrifibrocyte), and 28 (matrifibrocyte) after MI were subjected to RNA-seq. RNA-seq data were validated by principal component analysis (PCA) and Poisson distance analysis (**Figure 2AB**). Distinct transcriptomes were identified in cardiac fibroblasts of different states. The expression of a total of ∼20,000 genes was detected. Around 5,000 were differentially expressed between groups including both upregulated and downregulated genes (**Figure 2CD**). **Figure 2E** shows the expression of some representative genes which are divided into different groups based on their functions. The expression of proliferation genes peaked at day 3 after MI, which is in line with the peak proliferation rate of cardiac fibroblasts at this time point (**Figure 2E**). Stress fiber genes, myocardium ECM genes, cell migration genes were most enriched at both days 3 and 7 post-MI, corresponding to the myofibroblast state of cardiac fibroblasts (**Figure 2E**). Moreover, the expression of select cartilage genes was most enriched at day 28 post-MI, indicating the matrifibrocyte identity of cardiac fibroblasts in more stabilized infarcts (**Figure 2E**). Gene ontology (GO) analysis of genes that were upregulated in cardiac fibroblasts isolated from infarcts at different time points after MI as compared to cardiac fibroblasts isolated from uninjured hearts showed the enrichment of multiple biological processes. Some representative GO terms are listed in **Table 1**. The data is largely consistent with our previous publication (1).

**Table 1.** Genes of different functions were enriched in cardiac fibroblasts of different states. Genes that were upregulated in cardiac fibroblast isolated from infarct at different time points after MI as compared to cardiac fibroblasts from uninjured myocardium were subjected to GO enrichment analysis. Some most enriched GO terms and their enrichment p-values were shown in the table. D3, day 3; D7, day 7; W2, week 2; W4, week4.

|  | D3 | D7 | W2 | W4 |
| --- | --- | --- | --- | --- |
| GO_CELL_CYCLE | 2.27E-75 |  |  |  |
| GO_CYTOSKELETON_ORGANIZATION | 6.63E-38 |  |  |  |
| GO_REGULATION_OF_RESPONSE_TO_STRESS | 2.90E-33 |  |  |  |
| GO_CELL_MOTILITY | 2.31E-31 | 1.30E-59 | 1.95E-36 | 1.80E-45 |
| GO_APOPTOTIC_PROCESS | 3.18E-40 | 9.79E-39 | 2.09E-24 | 1.73E-24 |
| GO_SECRETION | 1.60E-49 | 2.46E-61 | 3.16E-33 | 4.74E-29 |
| GO_EXOCYTOSIS | 3.85E-51 | 2.98E-57 | 1.66E-21 |  |
| GO_DEFENSE_RESPONSE | 6.88E-52 | 5.51E-55 | 2.28E-28 |  |
| GO_BIOLOGICAL_ADHESION |  | 1.30E-59 | 1.76E-39 | 7.86E-47 |
| GO_EXTRACELLULAR_STRUCTURE_ORGANIZATION |  |  | 1.38E-36 | 1.32E-43 |
| GO_SKELETAL_SYSTEM_DEVELOPMENT |  |  | 1.68E-25 | 7.31E-24 |
| GO_OSSIFICATION |  |  | 3.07E-24 | 9.22E-25 |

**Figure 1.**
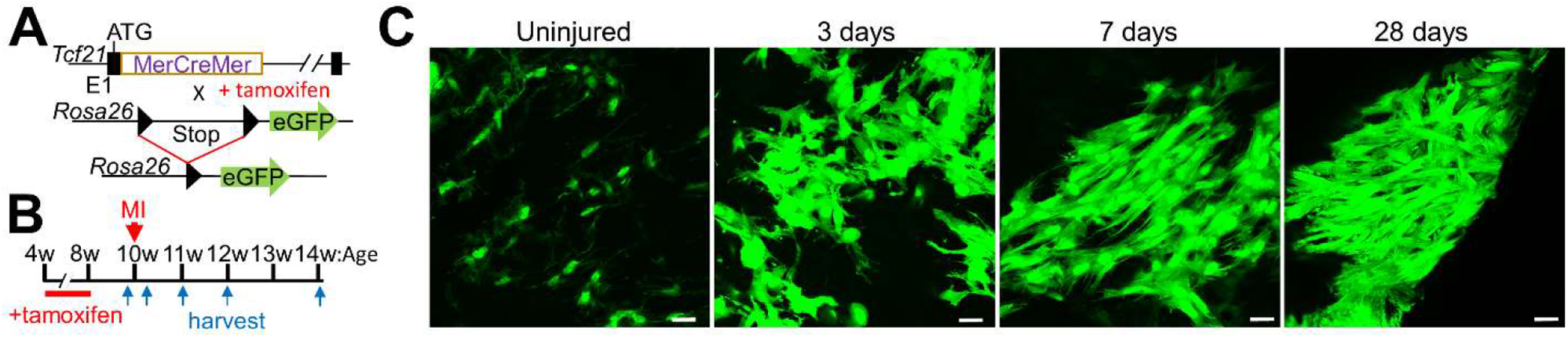
Lineage-tracing of cardiac fibroblast after MI. (**A**) Schematic of the *Tcf21*^*MCM/+*^;*R26*^*eGFP*^ mouse line. (**B**) Experimental scheme whereby *Tcf21*^*MCM/+*^;*R26*^*eGFP*^ mice were treated with TM, subjected to MI surgery, and then euthanized for heart sample collection at different time points after MI. (**C**). Whole-mount confocal microscopy images showing *Tcf21* lineage-traced cardiac fibroblasts (GFP^+^) in uninjured myocardium and infarcts at indicated time points after MI. Scale bar = 10 µm.

**Figure 2.**
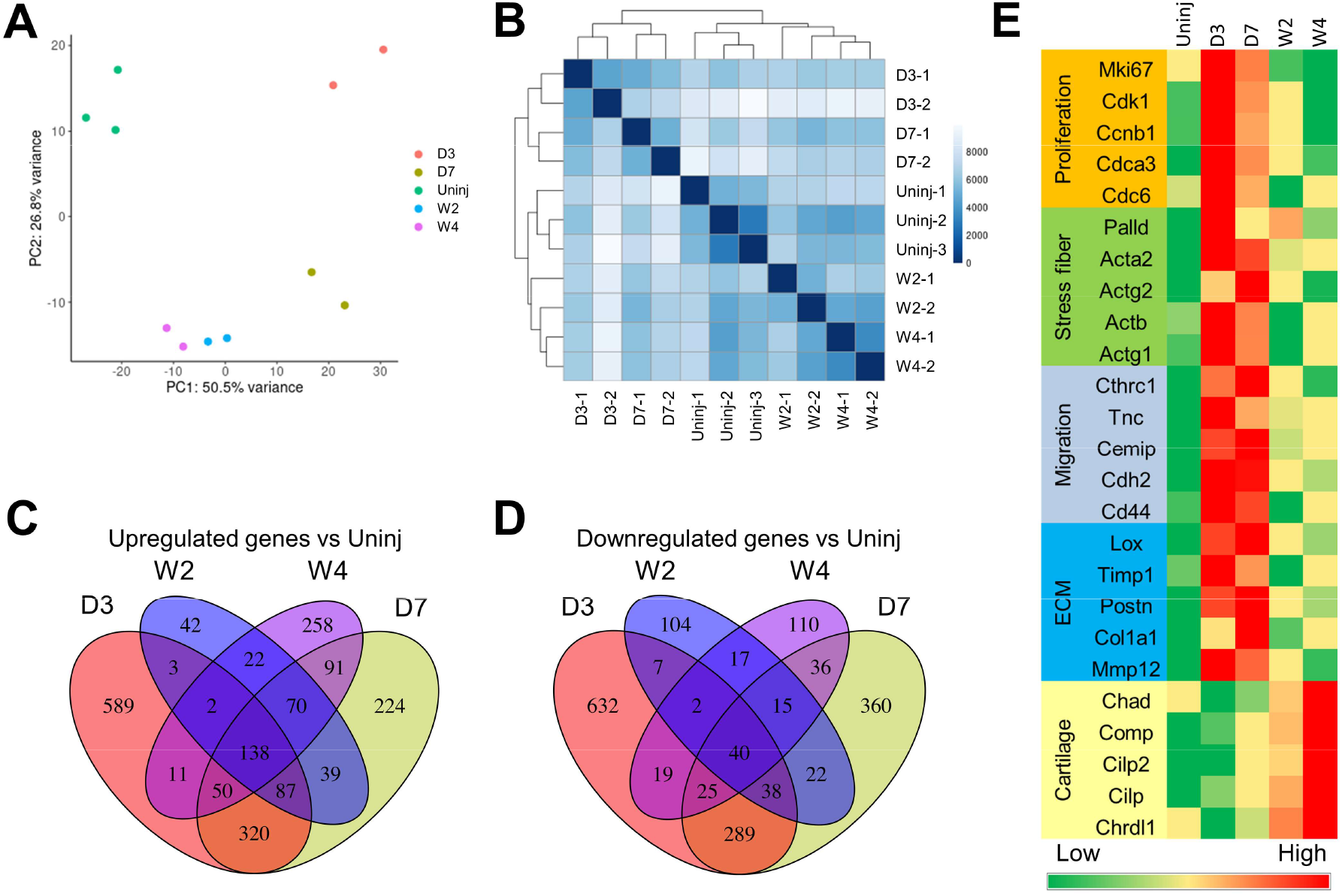
RNA-seq analysis of cardiac fibroblasts of different differentiation states. Tcf21 lineage-traced cardiac fibroblasts were sorted from uninjured myocardium and infarct at different time points after MI for RNA-seq analysis. (**A-B**) PCA analysis (A) and Poisson distance analysis (B) of RNA-seq data showing tight clustering of repeats of each group. (**C-D**) Venn diagram showing the overlaps between genes that were differentially expressed between *Tcf21* lineage–traced (EGFP+) fibroblasts isolated from the uninjured hearts and the other 4 groups of cardiac fibroblasts isolated from the MI region at different time points after injury. Genes that were upregulated compared to cardiac fibroblasts isolated from uninjured are shown in **C**. Downregulated genes are shown in **D**. (**E**) Heat map showing the normalized gene expression of some representative genes in different categories. Uninj, uninjured; D3, day 3; D7, day 7; W2, week 2; W4, week4.

### Accessible chromatin landscape in cardiac fibroblasts of different differentiation state

To investigate the chromatin accessibility in cardiac fibroblasts during their post-MI activation and differentiation, we performed ATAC-seq analysis on *Tcf21* lineage-traced cardiac fibroblasts isolated from the left ventricle of uninjured hearts and infarcts at day 3, 7, 14, and 28 after MI. Using our recently improved ATAC-seq protocol that is suitable for low cell numbers, we were able to obtain high-quality ATAC-seq libraries and data using 10,000 cells per sample. ATAC-seq peaks were highly enriched in the regions around the transcription start site (TSS) and around the transcription end site (TES) (**Figures 3AB**). In addition, ATAC-seq peaks were also highly enriched at CpG islands, suggesting that DNA methylation might play a role in chromatin remodeling during the activation and differentiation of cardiac fibroblasts (**Figure 3B**). An increasing number of studies have shown that repetitive elements such as interspersed elements (LINEs), short Interspersed elements (SINEs), long terminal repeats (LTR), and simple sequence repeats (SSRs) are prevalent in mammalian genomes and play some important roles in the regulation of chromatin accessibility and gene expression (22–27). However, we did not find significant enrichment of ATAC peaks in these repetitive elements or any difference in the enrichment of ATAC peaks in these repetitive elements among different sample groups, which suggests that repetitive elements are not the major regulator responsible for the changes in chromatin accessibility and gene expression during the post-MI activation and differentiation of cardiac fibroblasts (**Figure 3B**). PCA and Poisson distance analysis showed high sample consistency within each group and distinct chromatin accessibility profiles among different groups (**Figures 3CD**). Around 200,000 peaks were identified, among which 100,000 showed differential accessibility between cardiac fibroblasts of different groups (**Figures 3EF**).

**Figure 3.**
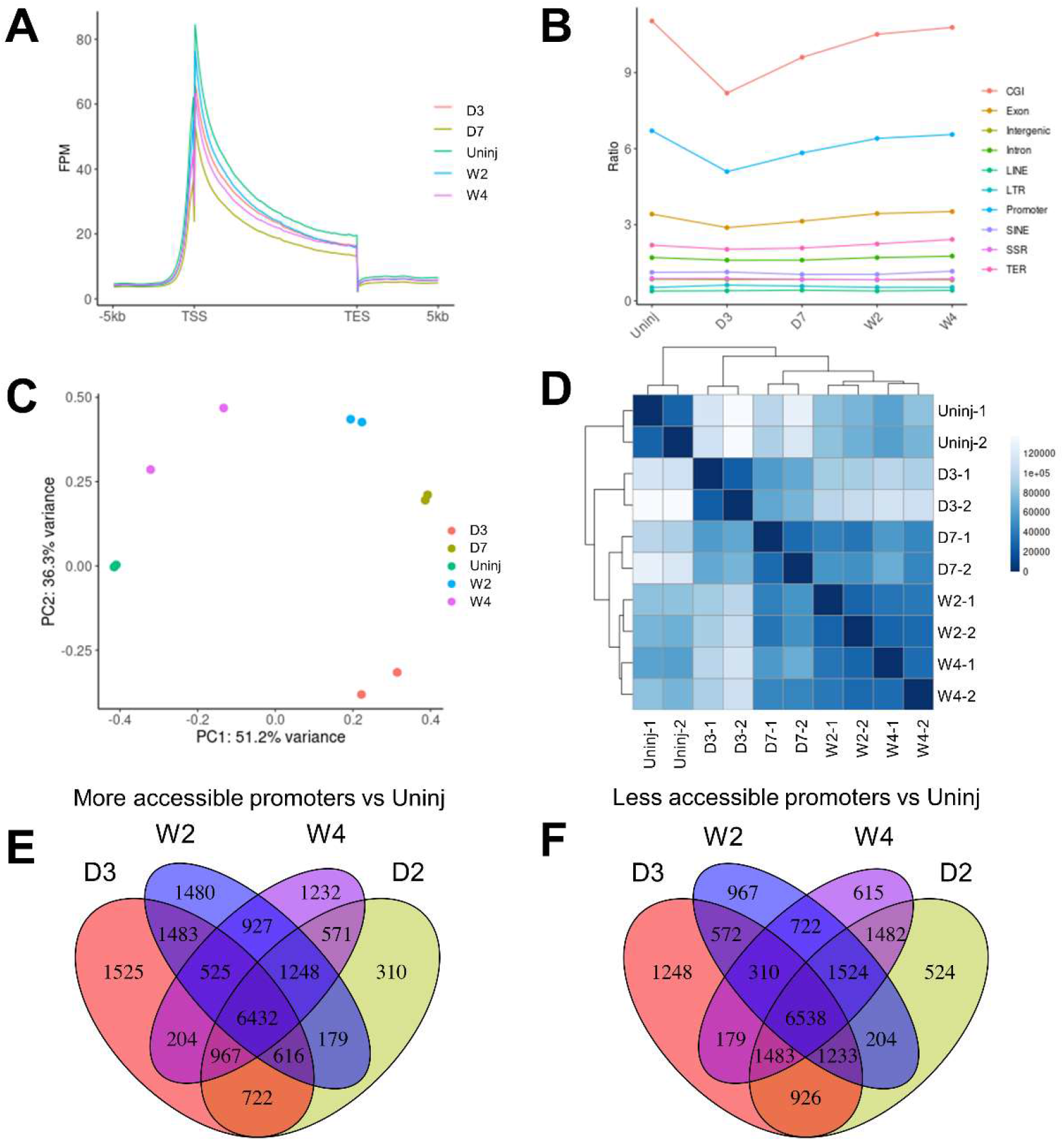
ATAC-seq analysis of cardiac fibroblasts of different differentiation states. Tcf21 lineage-traced cardiac fibroblasts were sorted from uninjured myocardium and infarct at different time points after MI for RNA-seq analysis. (**A**) Enrichment of ATAC-seq peaks around TSS and TES of genes. (**B**) Relative abundance of ATAC-seq peaks in different genomic features. (**C-D**) PCA analysis (C) and Poisson distance analysis (D) of ATAC-seq data showing tight clustering of repeats of each group. (**E-F**) Venn diagram showing the overlaps between ATAC-seq peaks that had different abundance between *Tcf21* lineage–traced (GFP+) fibroblasts isolated from the uninjured hearts and the other 4 groups of cardiac fibroblasts isolated from the MI region at different time points after injury. Peaks that were upregulated compared to cardiac fibroblasts isolated from uninjured are shown in **C**. Downregulated peaks are shown in **D**. Uninj, uninjured; D3, day 3; D7, day 7; W2, week 2; W4, week4.

### Differential gene expression between cardiac fibroblasts of different differentiation states is associated with dynamic chromatin accessibility

Chromatin accessibility of genes especially in the promoter regions may directly affect the transcription activity of genes. To study if promoter accessibility is correlated with gene expression in CFs, RNA-seq and ATAC-seq data were pulled together for an integrated analysis. It was found that the promoter accessibilities of genes expressed by at least one group of CFs were significantly higher than that of genes not expressed by any groups of cardiac fibroblasts (**Figure 4A**). An analysis of the 4957 genes that were differentially expressed between cardiac fibroblast groups shows that a positive correlation between promoter accessibility and transcription were identified in many differentially expressed genes (**Figure 4B**), such as *Tcf21* (a quiescent cardiac fibroblast marker), *Ki67* (a proliferation marker), *Acta2* (a myofibroblast marker), and *Comp* (a matrifibrocyte marker) (**Figure 4C-F**).

**Figure 4.**
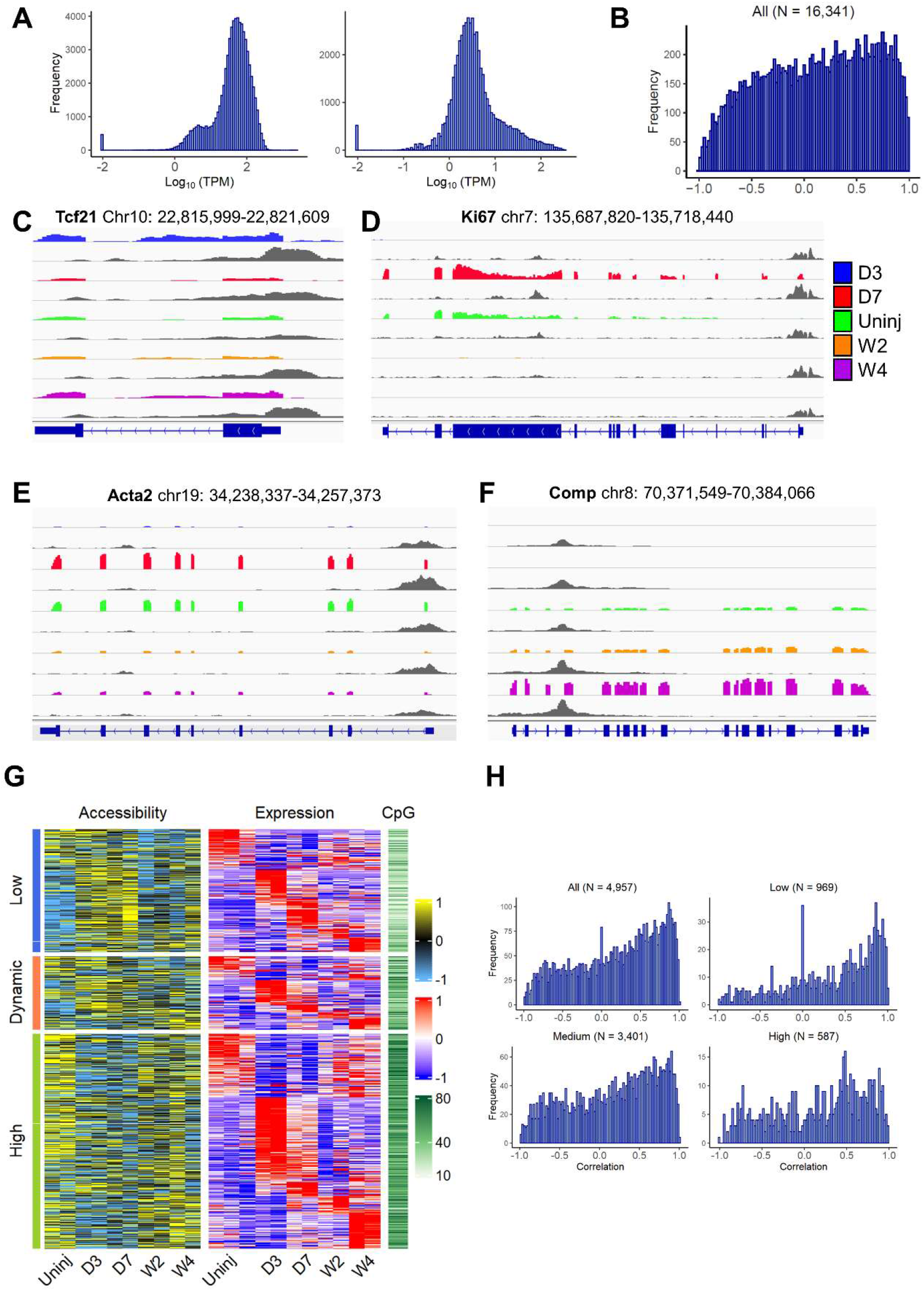
Differential gene expression and corresponding changes in promoter accessibility among cardiac fibroblasts of different differentiation states. (**A**) Relative promoter accessibility of genes that were expressed (left) or unexpressed (right) in cardiac fibroblasts measured by ATAC-seq. (**B**) Correlation between promoter accessibility and gene expression of all genes that were identified in RNA-seq and ATAC-seq analyses. (**C**-**F**) Integrated Genome Viewer (IGV) views of the RNA-seq data and ATAC-seq data of representative genes showing differential gene expression and promoter accessibility among *Tcf21* lineage-traced cardiac fibroblasts isolated from uninjured hearts and the MI region at different time points after injury. (**G**) Heat map showing the promoter accessibility, gene expression, and promoter CpG density of genes that were differentially expressed among cardiac fibroblasts isolated at different time points. Genes were divided into 3 groups based on their promoter accessibility (low, dynamic, and high). (**H**) Correlation between promoter accessibility and expression of all differentially expressed genes and genes that have low, medium, and high CpG densities. Uninj, uninjured; D3, day 3; D7, day 7; W2, week 2; W4, week4.

We then classify the detected promoters based on their accessibility. Accessibility data of each sample were standardized to have mean 0 and standard deviation 1 across promoters of genes included. Promoters received a standardized value that is no more than 0 in any of the samples were deemed to have constantly low accessibility. Those with values above 0 in all samples were deemed to have constantly high accessibility. The rest were deemed to have dynamic accessibility. The 4957 differentially expressed genes were then assigned to these 3 promoter accessibility groups. Interestingly, even though a strong promoter accessibility-transcription correlation was identified in many genes of all 3 promoter accessibility groups, a weaker correlation or lack of correlation was detected in some genes especially for those with high promoter accessibilities, suggesting that some other mechanisms besides chromatin accessibility may regulate the expression of genes that have generally more accessible promoters (**Figure 4G**). DNA methylation also regulates gene transcription activity. Increased DNA methylation may result in the recruitment of histone modification enzymes, which induces chromatin accessibility changes (28, 29). DNA methylation may also regulate gene expression through inhibiting TF binding (30, 31). We then looked at the CpG density in the promoters of different accessibility categories. Interestingly, genes with high promoter accessibility have the highest CpG density, and genes with low promoter accessibility have the lowest CpG density. These data suggest that DNA methylation may be a mechanism contributing to the low promoter accessibility-transcription correlation of some genes with constitutively high promoter accessibility (**Figure 4G**). Indeed, when the 4957 differentially expressed genes were divided into 3 groups based on the CpG densities in their promoter, a higher percentage of genes with low CpG density had stronger promoter accessibility-transcription compared to the other 2 groups, further suggesting the presence of gene transcription regulation by DNA methylation that is independent of chromatin accessibility (**Figure 4H**). Future study integrating DNA methylome analysis is required to reveal the crosstalk between DNA methylation and chromatin remodeling during the post-MI activation and differentiation of cardiac fibroblasts.

### Prediction of gene transcription changes by chromatin accessibility

The analysis performed above focused on the identification of corresponding changes in chromatin accessibility in genes that were differentially expressed between cardiac fibroblasts of different states. We then selected genes with differential promoter accessibility and those with differential accessibility in distal regions and study the changes in their transcription levels between different states of CFs. Consistently, among 3518 differentially accessible promoters, a strong positive correlation between promoter accessibility and transcription was detected in many genes (**Figure 5AB**). It is noteworthy that most promoters with differential accessibility belong to the low accessibility group (**Figure 5A**). Similarly, among these 3518 differentially accessible promoters, the strongest positive correlation was identified in promoters with low CpG density, further suggesting that high CpG density may be correlated with more active gene transcription regulation by DNA methylation which may regulate gene transcription without affecting promoter accessibility (**Figure 5B**). Enhancers located in distal regulatory regions also play important roles in gene transcription regulation (32). A total of 41713 distal ATAC-seq peaks that showed differential accessibility were identified. We then analyzed the accessibility of these ATAC-seq peaks and the expression of their most adjacent genes. Even though strong positive correlations between distal region accessibility and gene transcription were identified, the frequency of strong correlation was lower than that between promoter accessibility and gene transcription, which may be partially due to some incorrect assignments of ATAC-seq peaks to genes (**Figure 5CD**). Moreover, the majority of distal ATACseq peaks had low CpG density (**Figure 5D**), suggesting that DNA methylation may not play a major role in the regulation of the interaction between TFs and enhancers.

**Figure 5.**
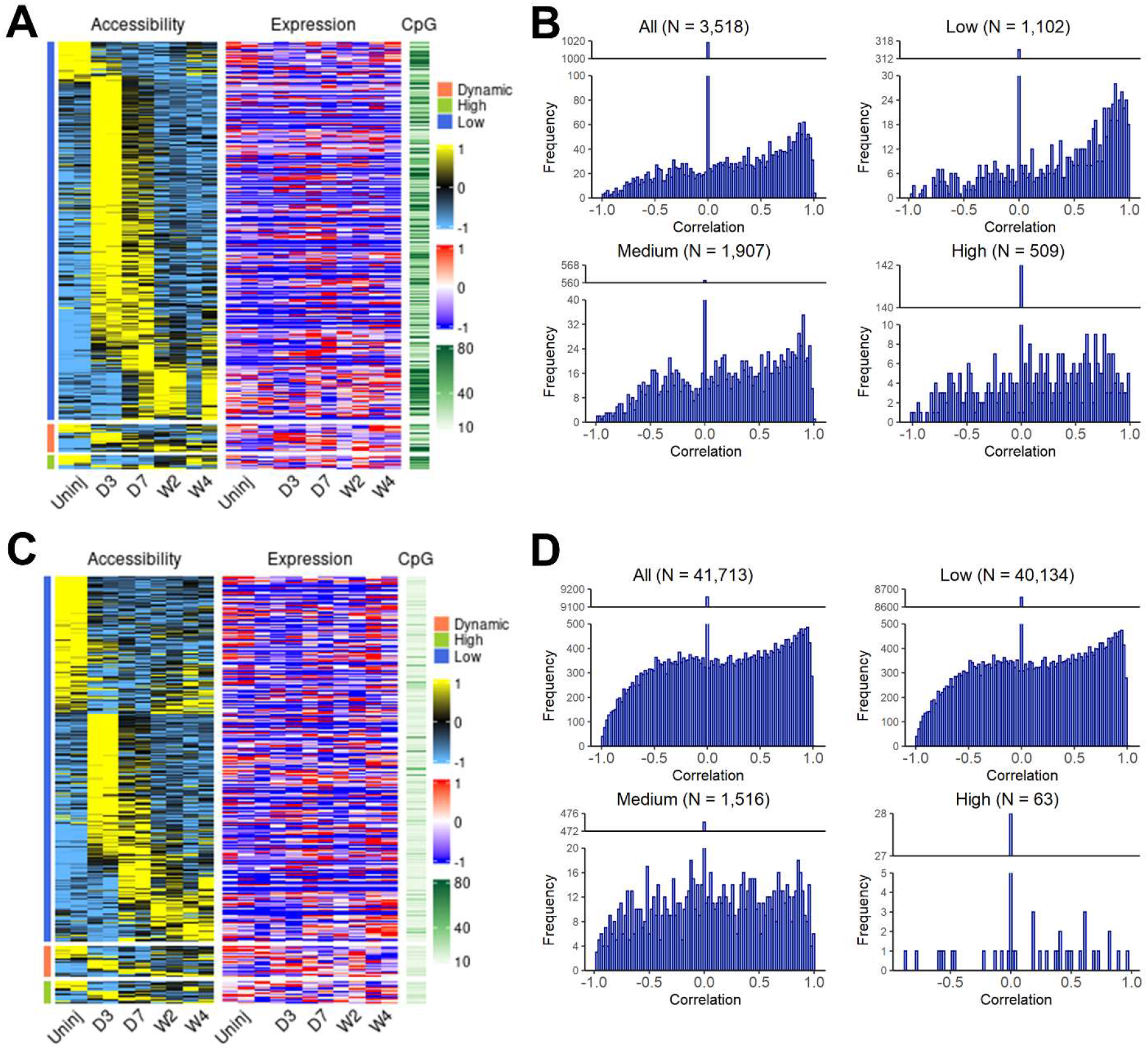
The correlation between differential accessibility in the promoter and distal regions and changes in the expression of corresponding genes among cardiac fibroblasts of different differentiation states. (**A**) Heat map showing the accessibility, expression level of corresponding genes, and CpG density of promoters that had differential accessibility among *Tcf21* lineage-traced cardiac fibroblasts isolated from uninjured and those isolated from MI region at different time points after injury. Genes were divided into 3 groups based on their expression (low, dynamic, and high). (**B**) Correlation between accessibility and expression of corresponding genes of all differentially accessible promoters and those of differentially accessible promoters that have low, medium, and high CpG densities. (**C**) Heat map showing the accessibilities, expression of corresponding genes, and CpG density of distal regions that had differential accessibility among *Tcf21* lineage-traced cardiac fibroblasts isolated from uninjured (Uninj) and the MI region 3 days (D3), 7 days (D7), 2 weeks (W2), or 4 weeks (W4) after injury. Genes were divided into 3 groups based on their expression (low, dynamic, and high). (**D**) Correlation between accessibility and expression of corresponding genes of all differentially accessible distal regions and those of differentially accessible distal regions that have low, medium, and high CpG densities. Uninj, uninjured; D3, day 3; D7, day 7; W2, week 2; W4, week4.

### Prediction of enhancers in cardiac fibroblast using ATAC-seq and RNA-seq data

Another feature of ATAC-seq other than measuring chromatin accessibility is that it can be used to predict TF binding sites. Unlike Chromatin Immunoprecipitation Sequencing (ChIP-seq) which is highly accurate but usually only reveals binding sites of a single TF per assay, ATAC-seq is able to predict the locations of many known TF binding motifs in the genome at the same time. Studies have found that enhancers play an important role in gene expression. Using the findMotifsGenome.pl from HOMER, 440 known motifs were identified, out of which 263 corresponding TFs had detectable expression in cardiac fibroblasts. The most enriched motifs including those for zinc finger proteins (e.g. Ctcf and Egr2), bZIP TFs (e.g. AP-1 family, C/EBP family, Nfil3), Nf1, Tead family (e.g. Tead2), EBF family (e.g. Ebf2), Stat family (e.g. Stat1), ETS family (e.g. ETS1), NF-κB family (e.g. Rela), MEF2 family (e.g. Mef2d), Runx family (e.g. Runx1), bHLH proteins (e.g. Tcf21), and ROR family (e.g. RORa). To identify motifs and corresponding TFs that are most likely responsible for the post-MI differentiations of cardiac fibroblasts, these motifs were classified into several groups based on their enrichment in different states of cardiac fibroblasts. The enrichment of Ctcf motif was found to be relatively unchanged across cardiac fibroblasts of all states, suggesting a constitutively active function of Ctcf in cardiac fibroblasts of all different states (**Figure 6A**). Interestingly, the enrichment of RORa motif temporarily dropped in myofibroblast, and a prolonged and progressive reduction in the enrichment of Tcf21 motifs was identified in both myofibroblasts and matrifibrocytes (**Figure 6B**). The enrichment of most of other motifs was found to be temporarily increased in myofibroblasts and then decreased in matrifibrocytes to a level around or below their level in quiescent cardiac fibroblasts (**Figure 6CD**). However, some exceptions were identified. In matrifibrocyte, a sustained elevation in the enrichment was identified for 4 motifs which are Tead2, Mef2s, Egr2, and Runx1 (**Figure 6E**). **Supplementary figure 1** shows the prevalence of these TF motifs in ATAC-seq peaks and background peaks, which suggests that the changes in the prevalence in target peaks and background peaks together mediated the differential enrichment of these motifs. An integrated analysis with RNA-seq data was then performed to predict the TFs that were likely responsible for the enrichment of these motifs. Corresponding changes in the expression were identified in some of these TFs (**Figure 6A-E**). Surprisingly, the expression of most bZIP TFs, which represent the largest TF group whose motifs were significantly enriched in myofibroblasts, dropped in myofibroblasts (**Figure 6B**). One exception is Fosl1 whose expression was upregulated in myofibroblasts (**Figure 6B**). Other TFs for which a corresponding change in expression was identified include Ctcf, RORa, Nf1, Stat1, Tead2, Runx1, and Tcf21. Given the fact that the motif enrichment and expression of Ctcf were not different among cardiac fibroblasts of different states, it is likely not responsible for the gene expression change during myofibroblast and matrifibrocyte differentiation. We then assigned distal ATAC-seq peaks to their most adjacent genes. An analysis was performed to study the prevalence of TF motifs in total distal ATAC-seq peaks and those assigned to genes that were upregulated in post-MI cardiac fibroblasts as compared to quiescent cardiac fibroblasts. Such an analysis revealed that these identified top enriched TF motifs in cardiac fibroblasts were more prevalent in ATAC-seq peaks adjacent to genes that were upregulated in cardiac fibroblasts post-MI than all identified ATAC-seq peaks except for Ctcf (**Figure 7A-E**), further suggesting that Ctcf does not attribute to the gene expression alteration in cardiac fibroblast after MI. Remarkably, the greatest difference was observed for Nf1 motif whose prevalence in post-MI upregulated genes was 2 times higher than that in total distal ATAC-seq peaks (**Figure 7C**). These analyses suggest that Fosl1, RORa, Stat1, Nf1, Tead2, Runx1, and Tcf21 may contribute to the post-MI activation and differentiations of cardiac fibroblast. We then identified the genes which were differentially expressed between cardiac fibroblasts of different states and possibly targeted by each of these TFs (corresponding motifs were identified in associated distal peaks). A comparison of differentially expressed genes targeted by individual TFs found that the largest overlapping was among Tcf21, Runx1, Tead2, Nf1, and Fosl1 (**Figure 8**), suggesting that a large number of genes that are responsible for the post-MI activation and differentiation of cardiac fibroblasts are co-regulated by multiple key TFs.

**Figure 6.**
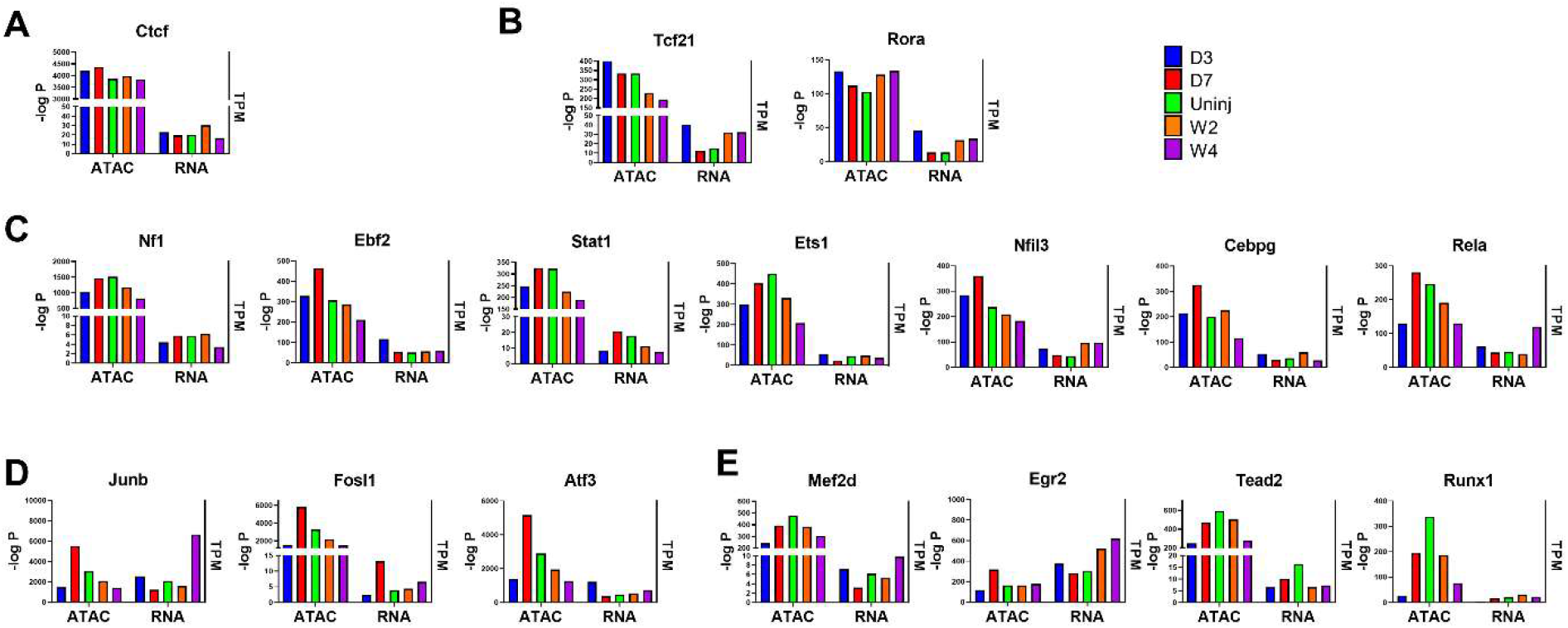
Identification of TF motifs in distal ATAC-seq peaks. The enrichment −log P values of some most enriched motifs and RNA expression levels of the corresponding TFs in *Tcf21* lineage-traced cardiac fibroblasts isolated from uninjured hearts and the MI region 3 days (D3), 7 days (D7), 2 weeks (W2), or 4 weeks (W4) after injury. Motifs were divided into uniformly enriched across all groups (A), more enriched in quiescent cardiac fibroblasts (B), temporarily more enriched in myofibroblasts (C-D), and more enriched in both myofibroblasts and matrifibrocytes.

**Figure 7.**
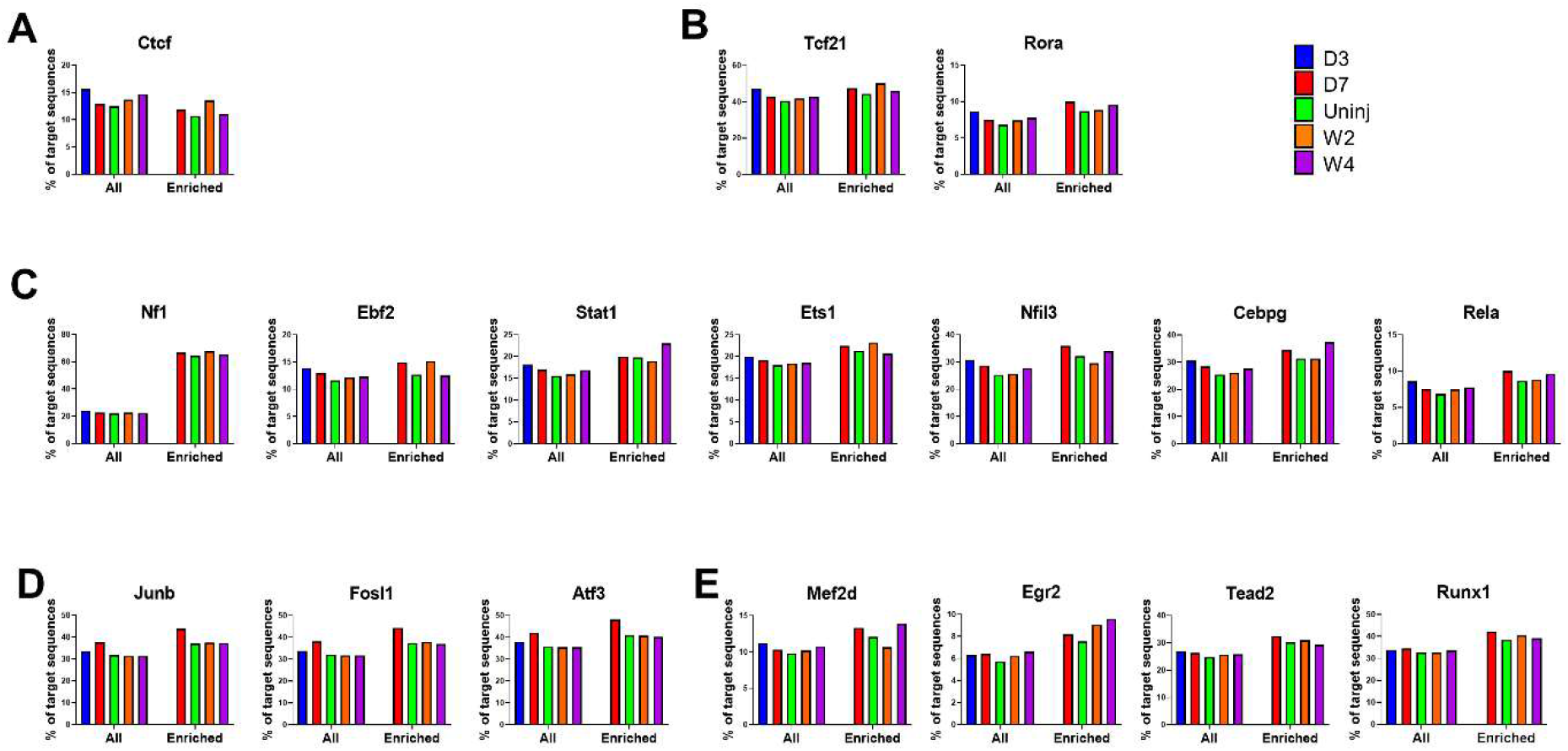
Increased enrichment of TF motifs in distal ATAC-seq peaks that are associated with genes upregulated in cardiac fibroblasts after MI. The prevalence (% of total peaks) of selected TF motifs in all distal ATAC-seq peaks (All) and in distal ATAC-seq peaks assigned to genes that were upregulated (Enriched) in cardiac fibroblasts at different time points after MI vs cardiac fibroblasts isolated from uninjured hearts. Motifs were divided into uniformly enriched across all groups (A), more enriched in quiescent cardiac fibroblasts (B), temporarily more enriched in myofibroblasts (C-D), and more enriched in both myofibroblasts and matrifibrocytes. Uninj, uninjured; D3, day 3; D7, day 7; W2, week 2; W4, week4.

**Figure 8.**
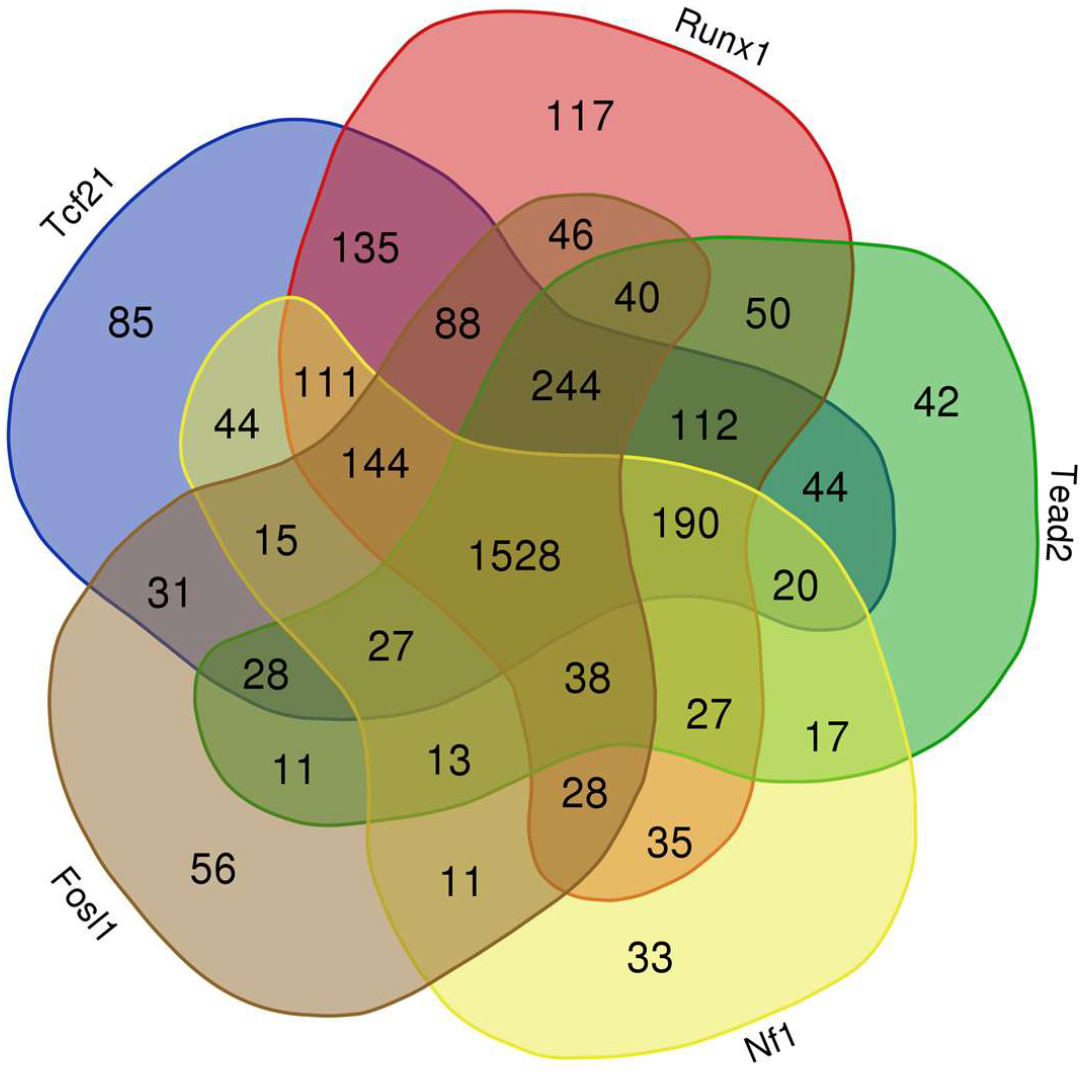
Overlaps between genes possibly targeted by individual TFs in the distal regions. Distal ATAC-seq peaks assigned to genes that were differentially expressed between cardiac fibroblasts of different states were analyzed for the prevalent of TF motifs. Venn diagram shows the overlaps between genes that are possibly targeted by individual TFs in the distal regions.

### Construction of gene regulatory network

We then identified the genes for which at least one motif of the 7 identified most significant motifs (Fosl1, RORa, Stat1, Nf1, Tead2, Runx1, and Tcf2) are located in ATAC-seq peaks ± 100 kb of their TSSs. The identified genes that were also in the top 500 genes that were most differentially expressed among cardiac fibroblasts of different states were selected for gene regulatory network construction (**Figure 9**). As expected, most genes in the gene regulatory network are regulated by more than 1 TFs. Most genes that are connected to Tcf21 and RORa appear to be negatively regulated by these 2 TFs (**Figure 9**). The requirement of Tcf21 in the epithelial-to-mesenchymal transition (EMT) during the cardiac fibroblast specification of epicardial cells has been reported (33). Mouse embryos lacking Tcf21 had reduced cardiac fibroblast number. Moreover, Tcf21 possibly also plays an important role in preventing the myofibroblast differentiation of cardiac fibroblasts as studies have found that the cells lacking Tcf21 express a higher level of αSMA and more readily differentiate into vascular smooth muscle cells (33–35). It was found that the expression of Tcf21 is regulated by C/EBPs whose binding motif is also enriched in cardiac fibroblast (36). Interestingly, retinoic acid signaling, which likely involves RORa, was also reported to play an important role during the prenatal differentiation of cardiac fibroblast (34), possibly through promoting the expression of Tcf21 (37). Indeed, a positive regulation of *Tcf21* expression by RORa was identified (Figure 9). Considering the stronger expression of Tcf21 and RORa in cardiac fibroblasts from uninjured hearts, they may have some important functions in maintaining the undifferentiated state of cardiac fibroblasts through inhibiting the expression of myofibroblast and matrifibrocyte marker genes. Surprisingly, even though both the expression and motif enrichment of Nf1 increased in cardiac fibroblasts after MI, Nf1 seems to function as a repressor during the post-MI activation and differentiation of cardiac fibroblasts (**Figure 9**). Similar to *Tcf21, Nf1* is also expressed in the epicardial precursors of cardiac fibroblasts (38). However, unlike *Tcf21*, the loss of *Nf1* resulted in increased EMT of epicardial cells and their proliferation. It is possible that the increased expression of *Nf1* in cardiac fibroblasts after MI is to downregulate the expression of genes that inhibit cardiac fibroblast activation and differentiation. Both positive and negative regulation of gene expression by Fosl2, Stat1, Tead2, and Runx1 were identified (**Figure 9**). Fosl1 is a member of the Fos family which is a part of a large TF family named AP-1 that also includes the Jun family and ATF family (39). Studies have found that members of AP-1 family regulate cell proliferation (40) and myofibroblast differentiation (41, 42). It is very interesting that despite the increased enrichment of AP-1 motifs in cardiac fibroblasts after MI, a reduction in the expression of most AP-1 family genes was observed except for Fosl1 which, however, was expressed at a rather low level (**Figure 6D**). To become functional, AP-1 subunits form heterodimers through the interaction of bZIP domains, which likely lead to their context-dependent function. It is possible certain mechanisms regulating the dimerization between specific AP-1 subunits are activated during the post-MI differentiation of cardiac fibroblasts after MI, which is likely more important than the expression levels of AP-1 subunits. A study comparing fetal and adult human cardiac fibroblasts identified Stat1 as an important TF responsible for the adult phenotype of cardiac fibroblasts (43). Loss of Stat1 in cardiac fibroblasts led to reduced cell size and collagen gene expression, and increased apoptosis. Thus, Stat1 may contribute to the increased ECM protein production by cardiac fibroblasts and their resistance to apoptosis after MI (1). Tead is a component of the Hippo pathway. Tead and its cofactor Yap1 were recently shown to promote both the fate determination of embryonic cardiac fibroblasts and the proliferation and myofibroblast differentiation of adult cardiac fibroblasts (44, 45). Runx1 has been shown to promote the proliferation of mesenchymal cells but delay their differentiation into myofibroblasts (46). A recent study showed that zebrafish with global Runx1 KO led to reduced cardiac fibrosis and a smaller number myofibroblast number after cardiac injury (47), suggesting an important function of Runx1 in cardiac fibroblasts and the development of cardiac fibrosis through balancing cardiac fibroblast proliferation and myofibroblast differentiation. Moreover, Runx1 is also important during the development of cartilage, a tissue shares some gene signature with matrifibrocytes (48). Runx1 is the only TF whose expression remains several folds higher in matrifibrocytes than in quiescent cardiac fibroblasts, accompanied by significantly higher motif enrichment (**Figure 6E**). Unlike other TFs that seem to coregulate the expression of genes, quite a number of genes seems to be solely regulated by Runx1 (**Figure 9**). Thus, other than its important roles in the early post-MI activation and myofibroblast differentiation of cardiac fibroblasts, Runx1 may also promote the matrifibrocyte differentiation in more stabilized infarct scar through regulating the expression of some unique genes.

**Figure 8.**
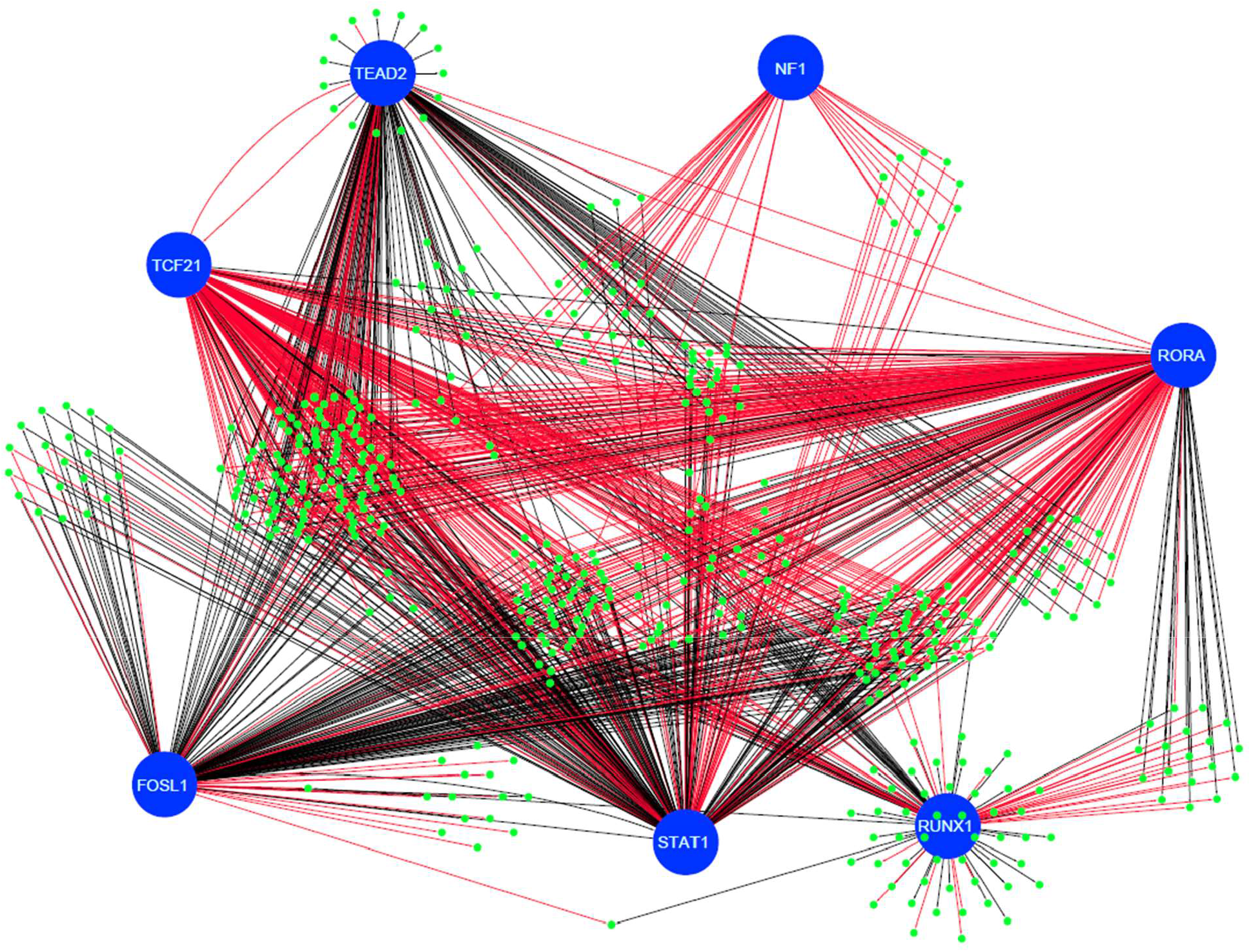
Gene regulatory network in cardiac fibroblasts. Top 500 most differentially expressed genes among cardiac fibroblasts of different states were analyzed for the presence of Fosl1, RORa, Stat1, Nf1, Tead2, Runx1, and Tcf2 motifs in ATAC-seq peaks ± 100 kb of their TSSs. Genes (green dots) with at least one identified motif and the 7 TFs (Fosl1, RORa, Stat1, Nf1, Tead2, Runx1, and Tcf2; blue circles) were selected for gene regulatory network construction. Red lines indicate negative regulation. Black lines indicate positive regulation.

## Conclusion

In conclusion, our transcriptome and open chromatin analysis showed that chromatin accessibility may play an important role in the post-MI activation and differentiation of cardiac fibroblasts. The integrated CpG density analysis also suggests that DNA methylation may regulate gene transcription through both regulating chromatin accessibility and a mechanism independent of chromatin remodeling. Finally, our motif analysis and gene regulatory network construction identified several potential TFs that are likely coordinated to mediate the sequential activation and differentiation of cardiac fibroblasts through programming the transcriptome of cardiac fibroblasts.

## Author contribution

X.F. and J.S. conceived the study; C.L., Q.L., H.M., L.W., and Y.L. collected samples. C.L., J.S., S.D., and R.L. analyzed data. C.L., J.S., J.F., and X.F. interpreted the data, assembled the results, and wrote the manuscript with inputs from all authors.

