## Supplemental figure 1 for "The landscape of accessible chromatin in quiescent and post-myocardial infarction cardiac fibroblasts"

p

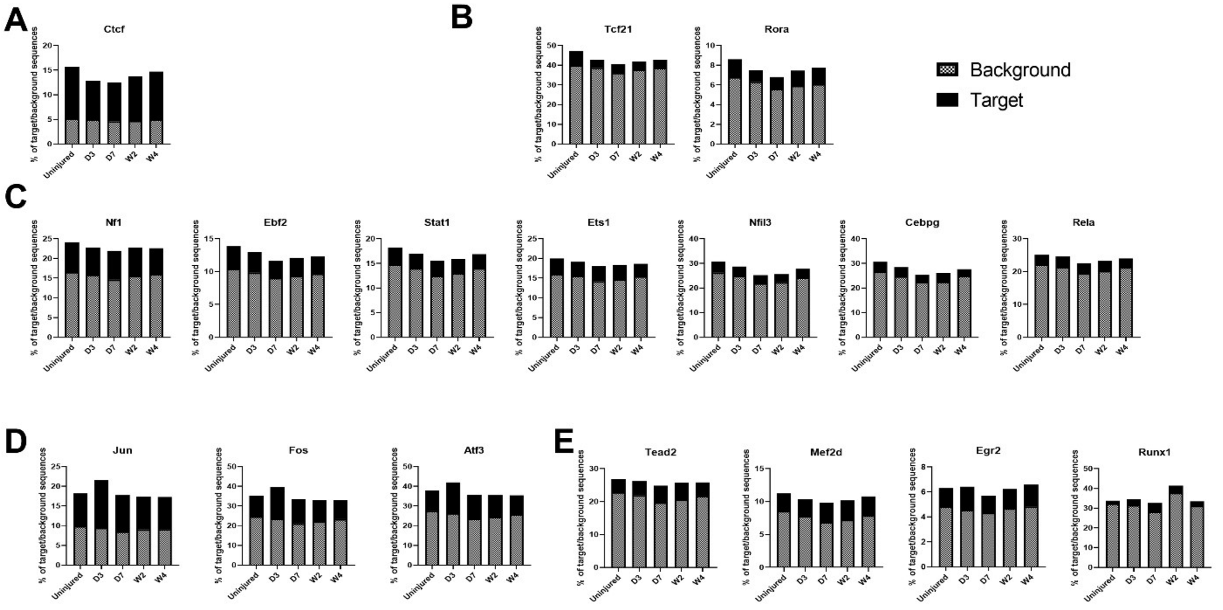


**Supplementary figure 1 (related to Figure 6.)**. The percentage of total distal ATAC-seq peaks and background peaks with selected individual TF motifs. Motifs were divided into uniformly enriched across all groups (A), more enriched in quiescent cardiac fibroblasts (B), temporarily more enriched in myofibroblasts (C-D), and more enriched in both myofibroblasts and matrifibrocytes. Uninj, uninjured; D3, day 3; D7, day 7; W2, week 2; W4, week4.
